# Maternal adiponectin restores cholinergic vasodilation in resistance vessels from adult offspring exposed to maternal obesity *in utero*

**DOI:** 10.1101/2024.09.21.612123

**Authors:** Molly Murphy, Sahand Fallahi, Jerad H. Dumolt, Julie A. Houck, Owen R. Vaughan, Lorna G. Moore, Colleen G. Julian, Fredrick J. Rosario, Theresa L. Powell, Thomas Jansson, Ramón A. Lorca

## Abstract

Maternal obesity increases the risk of cardiovascular and metabolic disease in the offspring both during childhood and adult life. Pregnant women and mice with obesity have lower circulating levels of adiponectin (ADN) compared to lean controls. ADN is an adipokine involved in regulating energy metabolism, vascular function, and placental function. We hypothesized that offspring of obese mice have impaired resistance artery function, which can be prevented by restoration of normal circulating ADN levels in obese dams during late pregnancy. Adult female mice were fed either control or obesogenic diet and mated with control diet-fed males. Control dams received a continuous infusion of phosphate saline buffer (PBS) during late pregnancy whereas obese females received either PBS or ADN. After weaning, offspring were fed a control diet. Mesenteric arteries (MsA) were dissected from adult offspring and mounted in a wire myograph or fixed for histology. MsA responses to vasoconstrictors (phenylephrine and endothelin-1) were not different between infusion groups. However, the vasodilatory responses to acetylcholine were reduced in offspring from obese dams as compared to control-fed dams. ADN supplementation during pregnancy restored the cholinergic vasodilatory responses of resistance vessels in offspring from obese dams. These observations suggest that normalizing circulating adiponectin levels in pregnancies complicated by obesity prevents in utero programming of vascular dysfunction in the offspring.

## INTRODUCTION

Obesity is the most common medical condition affecting pregnant women in developed countries (1). Children born to mothers with obesity are at an increased risk of being born large for gestational age, which can lead to birth complications such as shoulder dystocia and neonatal hypoglycemia (2). Maternal obesity during pregnancy is linked to an array of childhood and adult diseases in the offspring, in particular if born large for gestational age. These poor outcomes include coronary heart disease, stroke, type 2 diabetes, asthma, neurodevelopmental disorders, obesity, and epigenetic alterations in metabolic processes (3, 4, 5). The mechanisms underlying these effects are not fully understood, but it is believed that maternal obesity alters placental function and the fetal metabolic and endocrine environment, leading to changes in fetal growth, development, and programming of physiological systems (6).

While reducing the prevalence of obesity may be the most important long-term priority for population health, focus on this approach at the exclusion of other interventions may be counterproductive, especially due to the rising rates of obesity in women of reproductive age worldwide (7). Thus, research aimed at optimizing the health of offspring of obese mothers has high public health priority.

Maternal obesity has been shown to impair vascular function in the offspring (8, 9, 10) and has been associated with clinically relevant blood pressure elevation in children, raising concerns for future cardiovascular health (11). Impaired resistance artery vasodilation has been reported in offspring of obese rats, and epigenetic modulation of collagen and potassium channels, among other genes, has been proposed as the underlying mechanism (9).

Adiponectin (ADN) is a circulating adipokine secreted by adipose tissue that regulates metabolic processes such as glucose levels, lipid metabolism, and insulin sensitivity in several organs and tissues (12). Circulating levels of ADN increase during healthy pregnancies in the first and second trimesters and decrease in the third trimester when insulin sensitivity is reduced (13, 14). Plasma ADN levels are lower in obese pregnant women than in controls with normal body mass index (BMI) and negatively correlated with BMI (13, 15, 16, 17). Furthermore, ADN supplementation in obese pregnant mice has been associated with enhanced placental function, reduced fetal weight, and improved cardiac function and glucose metabolism in the offspring compared to obese untreated dams (15, 18, 19). However, the role of ADN supplementation in preventing vascular dysfunction in offspring of mothers with obesity has not been explored.

Here, we tested the hypothesis that offspring of obese mice have impaired resistance artery function, which can be prevented by restoration of normal circulating ADN levels in obese dams during late pregnancy. We isolated resistance vessels to study global vascular effects of maternal obesity and ADN supplementation in adult offspring born to control or obese dams infused with ADN or vehicle during pregnancy. Our results highlight targeting maternal ADN signaling, for instance with ADN receptor antagonist, as a promising therapeutic approach to prevent vascular dysfunction in adult offspring exposed to maternal obesity in utero.

## METHODS

### Mice

All procedures were approved by the Institutional Animal Care and Use Committee of the University of Colorado (Protocol # 00320) and conducted in accordance with NIH guidelines. The mouse model has been described in detail previously (15, 18, 19, 20) and the study design is presented in **Figure 1**. Briefly, 12-week-old C57/BL6J female mice (Jackson Laboratory), proven breeders (one previous litter), were randomly assigned to either a control diet (CON, D12489, Research Diets, 10% calories from fat, n = 5) or an obesogenic diet (OB, Western Diet D12089B, Research Diets, 40% calories from fat plus 20% sucrose solution *ad libitum*, n = 6). Females were mated with CON-fed males once animals in the OB group had gained 25% above of their initial body weight. A mini osmotic pump (Alzet 1003D, Alza Corporation) was subcutaneously implanted in isoflurane-anesthetized dams at embryonic day (E) 14.5. CON dams received a continuous infusion of sterile phosphate-buffered saline (PBS). OB animals were randomly allocated to receive either PBS (n = 3) or mouse recombinant full-length ADN from E14.5 through E18.5 (n = 3, ALX-522–059; Enzo Life Sciences, 0.62 μg*g of body weight^−1^*day^−1^), as the osmotic pumps cease functioning after 4 days. This infusion rate has been reported to increase circulating levels of ADN in pregnant OB mice to the levels observed in normal weight (CON) dams (15). Dams delivered spontaneously and were maintained on their respective diets throughout lactation. Offspring were weaned 28 days after birth, maintained on standard chow, and housed in same-sex groups from multiple litters. Study groups were divided into 1) offspring of CON diet-fed dams infused with vehicle (CON+PBS), 2) offspring of OB dams infused with vehicle (OB+PBS), and 3) offspring of OB dams infused with ADN (OB+ADN). Seven-to 9-month-old offspring of each group were euthanized by CO_2_ inhalation followed by cervical dislocation and mesentery dissection. *N* values are individual offspring mice, and because male and female offspring did not differ in vascular responses (data not shown) they were grouped together.

**Figure 1.**
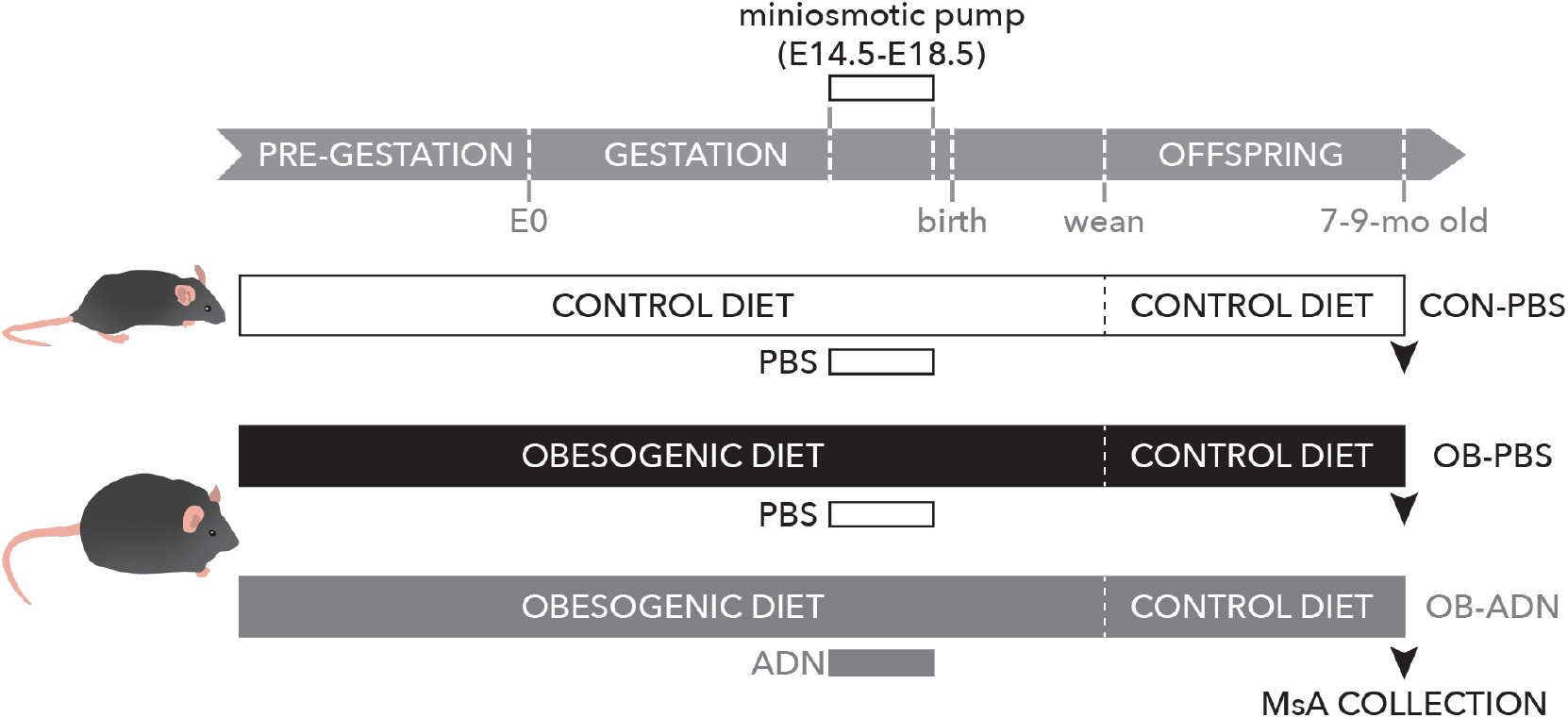
Study design. Females were fed a control (CON) or obesogenic (OB) diet pre-gestationally, throughout gestation and until pups were weaned. At embryonic day (E) 14.5, a mini osmotic pump was implanted in pregnant mice and phosphate saline buffer (PBS) or adiponectin (ADN) was infused. After weaning, pups were fed a CON diet until 7-9 months old when they were euthanized and MsA collected for functional studies.

### Mesenteric Artery Dissection and Wire Myography

Whole mesentery including intestines and blood vessels was isolated and placed in ice-cold PBS (Thermo Fisher Scientific) as previously reported (21). Second-to third-order mesenteric arteries (MsA) were dissected under a microscope in ice-cold PBS, and adipose and connective tissue surrounding the vessels were removed. After being cleaned and cut into segments (∼2 mm), the arteries were mounted in a small vessel wire myograph (Multi Wire Myograph System 610M, DMT-USA) and normalized in a warmed and oxygenated (95% O_2_/5% CO_2_) Krebs buffer (118 mM NaCl from Sigma Aldrich, 4.7 mM KCl, 2.5 mM CaCl_2_, 1.2 mM MgSO_4_, 1.2 mM KH_2_PO_4_, 25 mM NaHCO_3_, and 11 mM D-glucose, all from Thermo Fisher Scientific). Vessels were allowed to equilibrate in the chambers for at least 45 min after normalization. MsA were constricted with 120 mM KCl to assess viability. After washing, the vessels were exposed to increasing concentrations of the vasoconstrictor endothelin-1 (ET-1, 0.01 – 100 nM) or phenylephrine (PE, 0.1 nM – 100 µM, Sigma-Aldrich). MsA were then pre-constricted with 10 µM PE for 20 min and increasing concentrations of vasodilators were applied to the chambers; acetylcholine (ACh, 0.1nM – 100 µM, Acros Organics), bradykinin (BK, 0.1 nM – 1 µM; Anaspec), sodium nitroprusside (SNP, 0.1 nM – 100 µM; Sigma-Aldrich) or A769662, an AMP-activated protein kinase activator, (0.1 µM – 100 µM; Selleck Chemicals). To assess the contribution of endothelial signaling to ACh vasodilation, MsA were pre-incubated with the endothelial nitric oxide synthase inhibitor L-N^G^-nitroarginine methyl ester (L-NAME, 10 µM; Sigma-Aldrich) or L-NAME (10 µM) plus the cyclooxygenase inhibitor indomethacin (INDO, 10 µM; Sigma-Aldrich) for 20 min before PE-constriction and the application of ACh.

### Data Analysis

Offspring weights were measured at weaning and adult weight at euthanasia, and age at the time of euthanasia was calculated. The tension developed by ET-1 or PE was normalized to that elicited by 120 mM KCl (K_max_), whereas the vasodilatory responses were calculated as percentage of 10 µM PE constriction. Half-maximal effective concentration (EC_50_), maximal force (E_max_) and area under the curve (AUC) were measured by Graph Pad 9 software. Data from each group were tested for normality (Kolmogorov-Smirnov) and compared by unpaired one-way ANOVA or Kruskal-Wallis test, as appropriate, followed by multiple comparisons post hoc test. Offspring sex was compared by Fisher’s test. Significance was set at p < 0.05.

## RESULTS

### Offspring characteristics

Although the dams of the OB+PBS and OB+ADN groups remained obese (data not shown), offspring average body weight at weaning and adult body weight at time of euthanasia were similar between groups (**Table1**). The percentage of female offspring was also similar between groups (**Table 1**). The offspring from OB+ADN were a month or less than a month older than CON+PBS and OB+PBS groups, respectively (**Table 1**).

**Table 1.** Offspring characteristics.

| **Characteristic** | **CON+PBS** | **OB+PBS** | **OB+ADN** | ***p*-value** |
| --- | --- | --- | --- | --- |
| Number of litters [total number of offspring] | 5 [21] | 3 [11] | 3 [10] | N/A |
| Weight at weaning (g) | 11.6 ± 0.6 | 13.0 ± 0.4 | 13.6 ± 0.8 | 0.0872 |
| Age at euthanasia (months) | 7.9 ± 0.1 | 8.2 ± 0.2 | 8.9 ± 0.1*<sup>#</sup> | 0.0003 |
| Weight at euthanasia (g) | 29.4 ± 1.8 | 32.3 ± 2.0 | 35.7 ± 2.6 | 0.1345 |
| Female Sex (%) | 47.6 | 27.3 | 20 | 0.3122 |

Values are mean ± SEM. All values are individual mice. *p*-values were calculated via one-way ANOVA, Kruskal-Wallis or Fisher's test. \* *p* < 0.01 vs. CON+PBS, <sup>#</sup> *p* < 0.05 vs. OB+PBS.

### Vasoconstrictor and vasodilator responses in offspring MsA

We observed similar constrictions to both PE or ET-1 in MsA from all three groups (**Figure 2**), and no differences in EC_50_ or E_max_ values between groups (**Table 2**). The vasodilatory response to ACh was greatest in the CON+PBS and OB+ADN groups and decreased in the OB+PBS group (p = 0.0008 and p = 0.0325 compared to CON+PBS and OB+ADN, respectively) (**Figure 3A and 3B**). The vasodilatory responses to BK were not different between the three groups (**Figures 3C and 3D**).

**Table 2.**
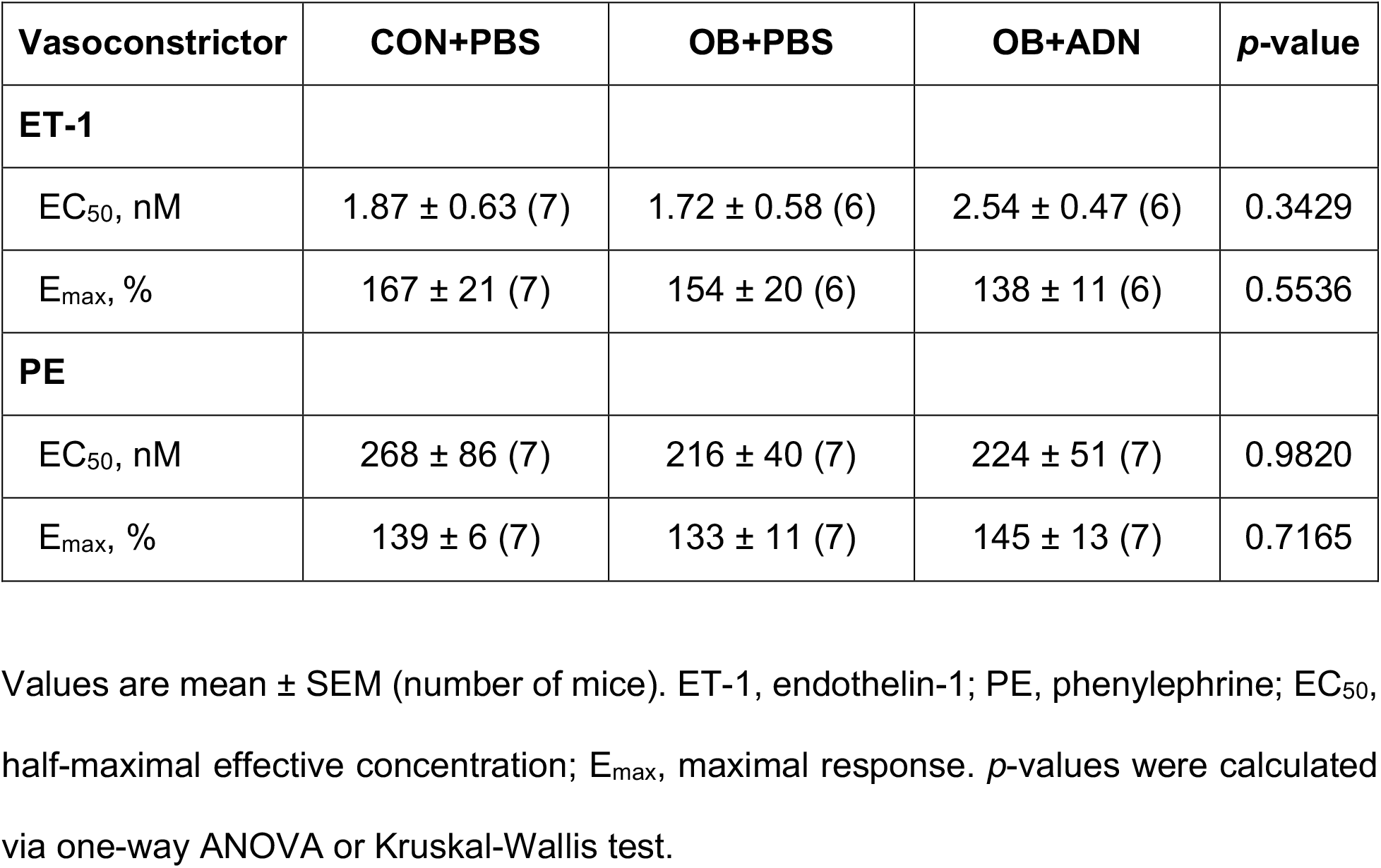
Vasoconstrictor responses in MsA from different groups.

| **Vasoconstrictor** | **CON+PBS** | **OB+PBS** | **OB+ADN** | ***p*-value** |
| --- | --- | --- | --- | --- |
| **ET-1** |  |  |  |  |
| EC<sub>50</sub>, nM | 1.87 ± 0.63 (7) | 1.72 ± 0.58 (6) | 2.54 ± 0.47 (6) | 0.3429 |
| E<sub>max</sub>, % | 167 ± 21 (7) | 154 ± 20 (6) | 138 ± 11 (6) | 0.5536 |
| **PE** |  |  |  |  |
| EC<sub>50</sub>, nM | 268 ± 86 (7) | 216 ± 40 (7) | 224 ± 51 (7) | 0.9820 |
| E<sub>max</sub>, % | 139 ± 6 (7) | 133 ± 11 (7) | 145 ± 13 (7) | 0.7165 |

Values are mean ± SEM (number of mice). ET-1, endothelin-1; PE, phenylephrine; EC<sub>50</sub>, half-maximal effective concentration; E<sub>max</sub>, maximal response. *p*-values were calculated via one-way ANOVA or Kruskal-Wallis test.

**Figure 2.**
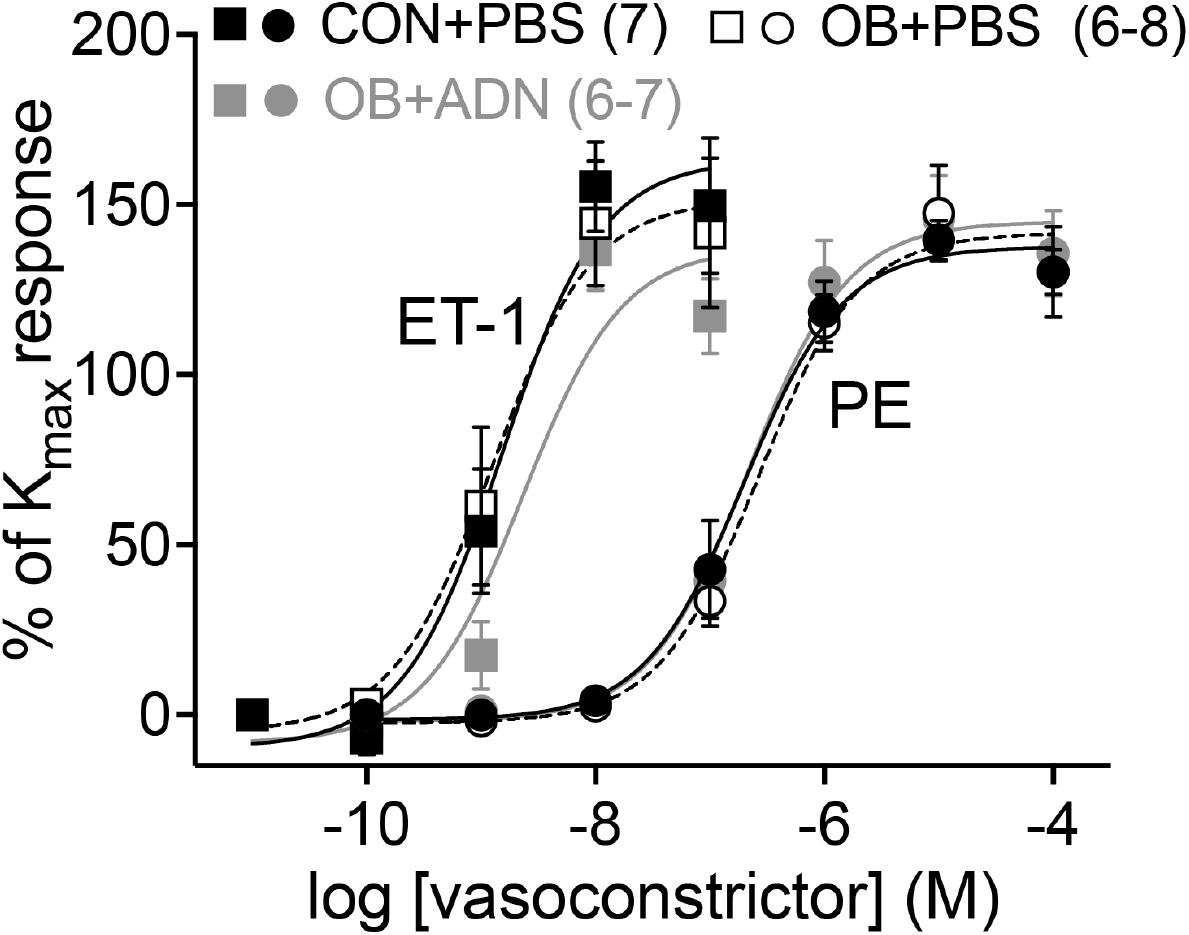
Mesenteric artery (MsA) responses to vasoconstrictors. Concentration-response curves of endothelin-1 (ET-1, squares) or phenylephrine (PE, circles) in MsA of offspring from control dams (CON+PBS, closed symbols), obese dams (OB+PBS, open symbols) or obese dams supplemented with ADN (OB+ADN, gray symbols), presented as percentage of maximal KCl response (K_max_). Numbers in parentheses are the number of mice, symbols are mean values ± SEM.

**Figure 3.**
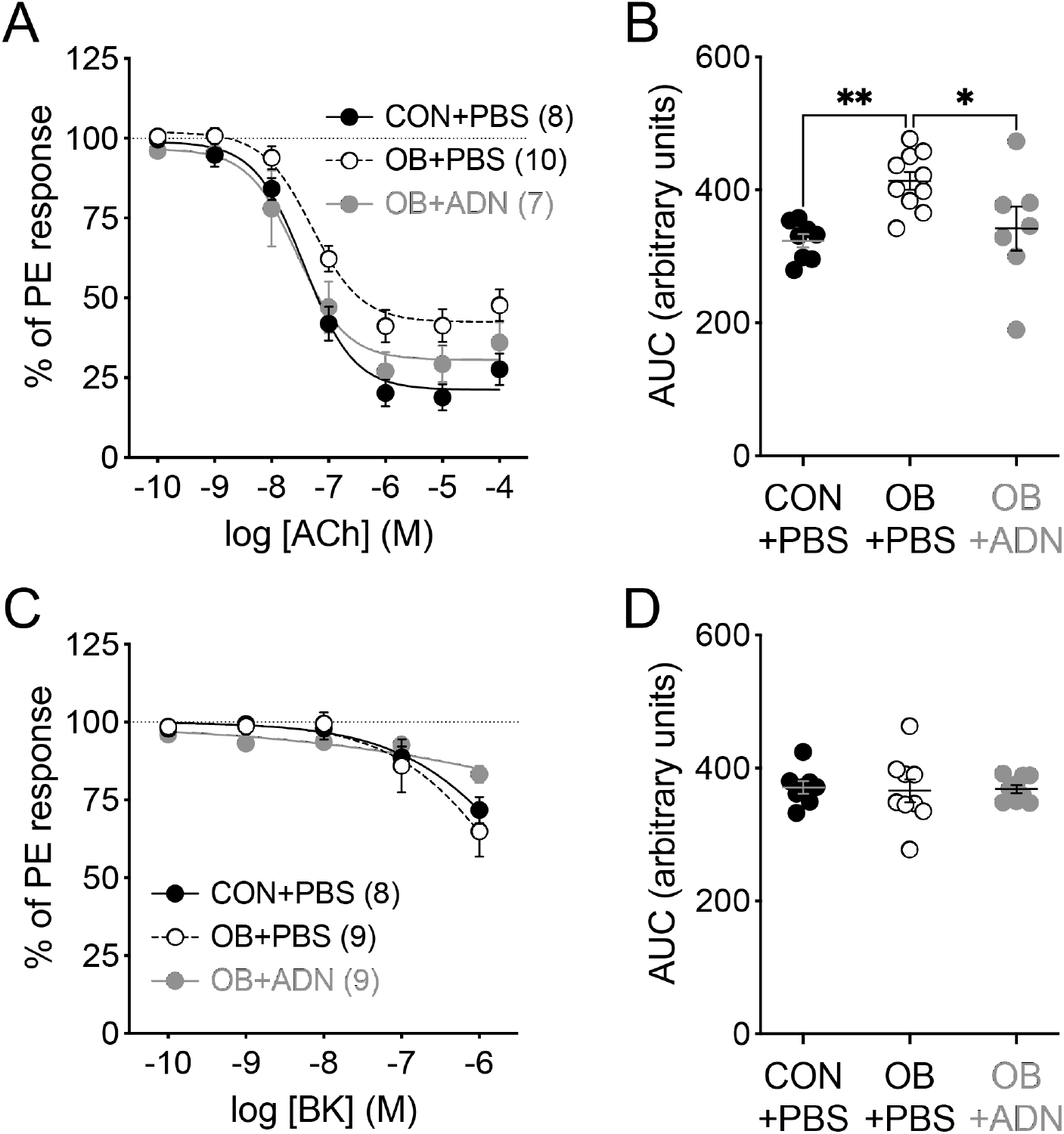
Mesenteric artery (MsA) responses to vasodilators. Concentration-response curves of acetylcholine (ACh, **A**) or bradykinin (BK, **C**) in MsA of offspring from control dams (CON+PBS, closed symbols), obese dams (OB+PBS, open symbols) or obese dams supplemented with ADN (OB+ADN, gray symbols), presented as percentage of 10 µM phenylephrine (PE) response. Numbers in parentheses are the number of mice, symbols are mean values ± SEM. **B** and **D** show the area under the curve (AUC) analysis of curves in A and C, respectively. * *p* < 0.05 and ** *p* < 0.01 by one-way ANOVA.

### Endothelial mechanisms of ACh-dependent vasodilation

In the CON+PBS offspring, the vasodilatory response to ACh was reduced in the presence of L-NAME (p = 0.0346), and further decreased in the presence of L-NAME and INDO (p < 0.01 and p = 0.0281 compared to untreated and L-NAME, respectively) (**Figure 4A and 4B**), suggesting that both signaling pathways are activated by ACh. However, in the OB+PBS offspring, the vasodilatory response to ACh was slightly, although not significantly, decreased by L-NAME (p = 0.075 compared to untreated), and a further, modest reduction was observed with L-NAME+INDO (p = 0.0424 compared to untreated) (**Figure 4C and 4D**), suggesting that the reduction in ACh responses found in the OB+PBS group may be due to a diminished nitric oxide signaling. In the OB+ADN offspring, the vasodilatory response to ACh did not differ from untreated in the presence of L-NAME or L-NAME+INDO (**Figure 4E and 4F**).

**Figure 4.**
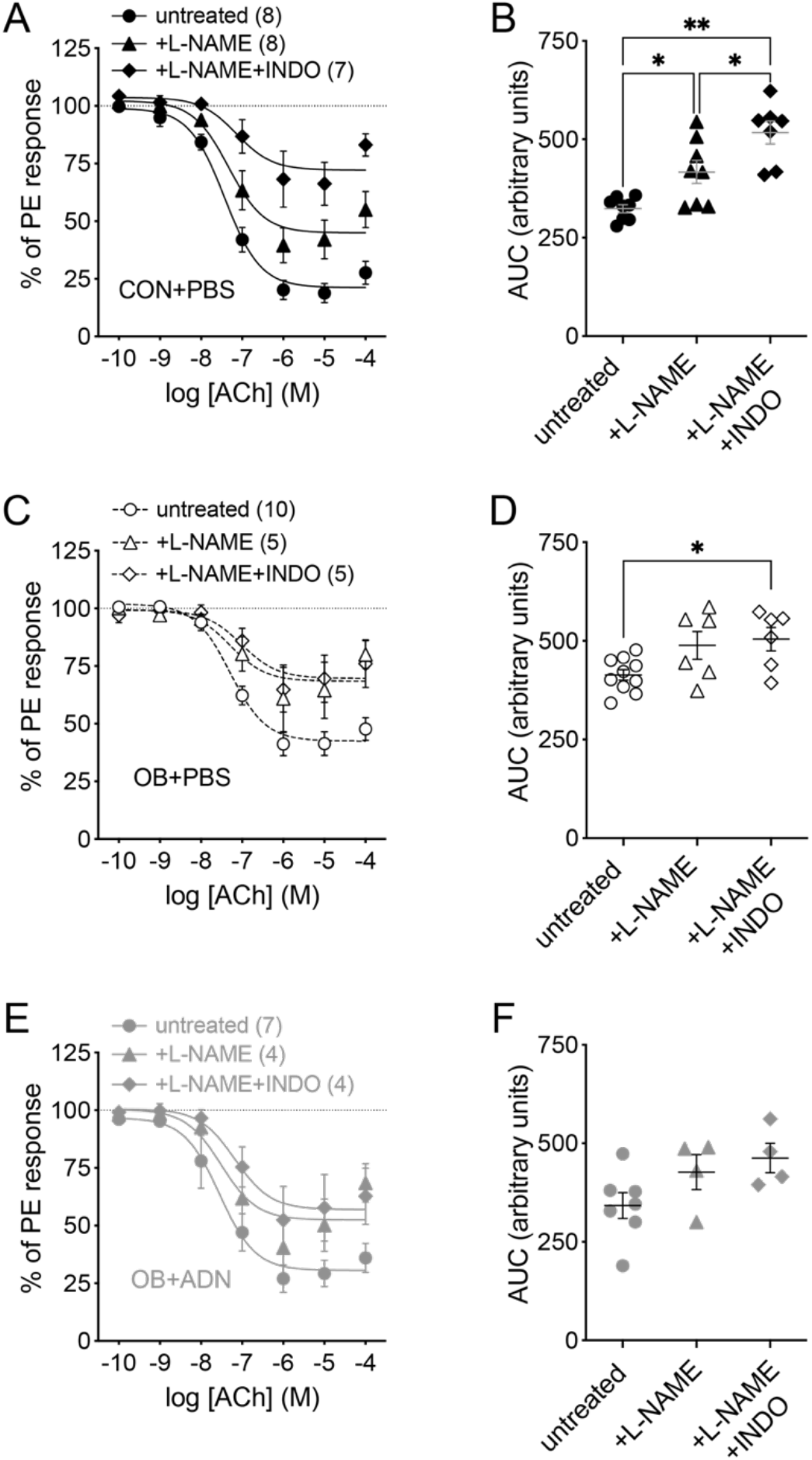
Mechanisms of mesenteric artery (MsA) responses to acetylcholine. Concentration-response curves of acetylcholine (ACh) in MsA of offspring from control dams (CON+PBS, closed symbols, **A** and **B**), obese dams (OB+PBS, open symbols, **C** and **D**) or obese dams supplemented with adiponectin (OB+ADN, gray symbols, **E** and **F**), in the absence (untreated, circles) or presence of 10 µM L-NAME (triangles) or 10 µM L-NAME + 10 µM indomethacin (INDO, diamonds). Numbers in parentheses are the number of mice. Symbols in **A, C** and **E** are mean values ± SEM presented as percentage of 10 µM phenylephrine (PE) response; symbols in **B, D** and **F** are individual values ± SEM presented as the area under the curve (AUC) of curves in **A, C** and **D**, respectively. * *p* < 0.05 and ** *p* < 0.01 by one-way ANOVA.

### Other vasodilatory responses in offspring MsA

The vasodilatory response to SNP was not different between CON+PBS and OB+PBS, whereas the concentration-response curve was shifted to the right in the OB+ADN group (**Figure 5A**). Area under the curve analysis shows a decrease in the vasodilatory responses in OB+ADN compared to CON+PBS (p = 0.0283) and OB+PBS groups (p = 0.0221) (**Figure 5B**). The vasodilatory response to A769662 did not differ between the 3 groups (**Figure 5C and 5D**).

**Figure 5.**
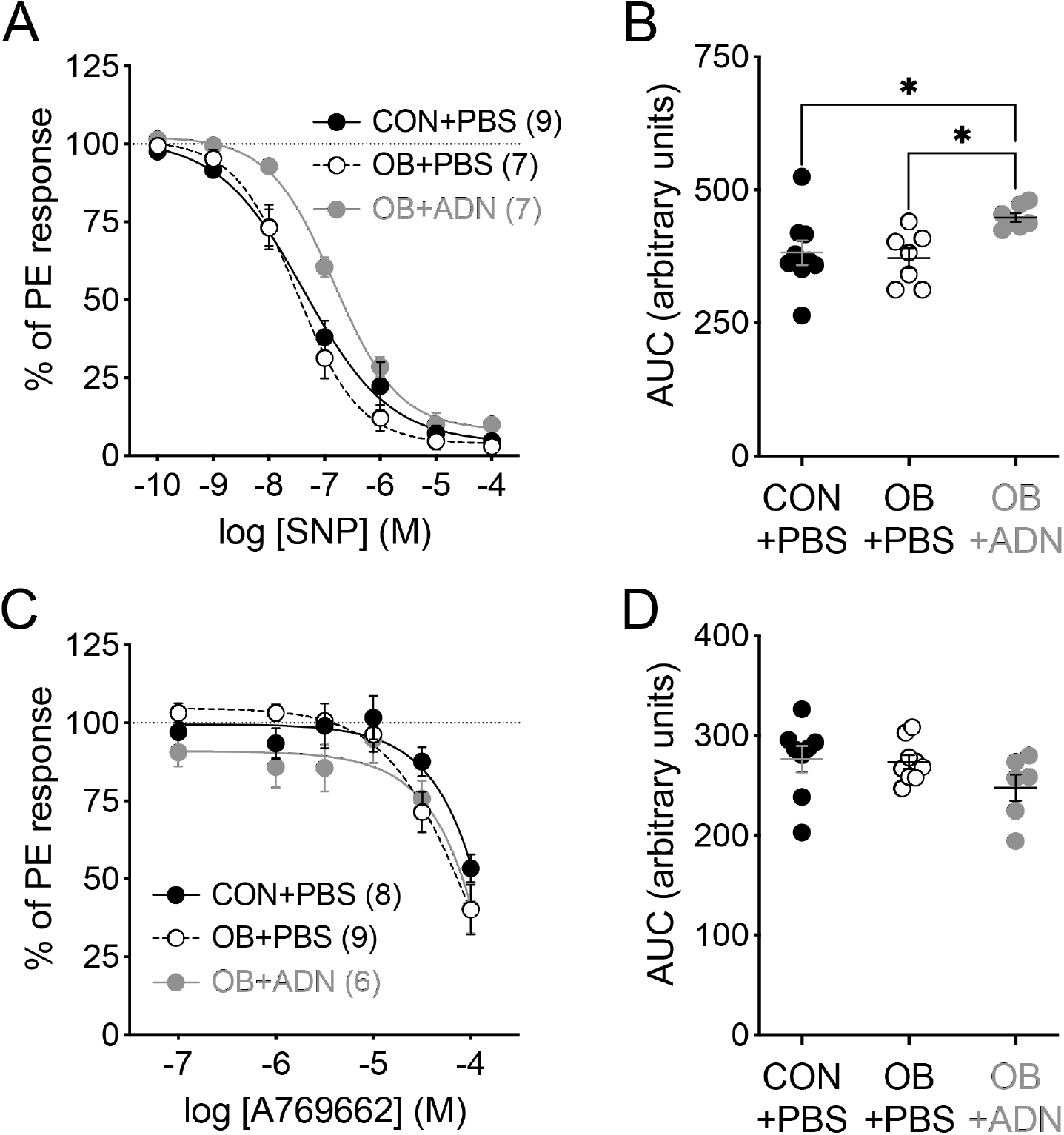
Responses of mesenteric arteries (MsA) to a nitric oxide donor and an AMPK activator. Concentration-response curves of sodium nitroprusside (SNP, **A**) or A769662 (**C**) in MsA of offspring from control dams (CON+PBS, closed symbols), obese dams (OB+PBS, open symbols) or obese dams supplemented with ADN (OB+ADN, gray symbols), presented as percentage of 10 µM phenylephrine (PE) response. Numbers in parentheses are the number of mice, symbols are mean values ± SEM. **B** and **D** show the area under the curve (AUC) analysis of curves in A and C, respectively. * *p* < 0.05 by one-way ANOVA.

## DISCUSSION

In this study, we observed that maternal obesity impairs vasodilation in resistance arteries of adult offspring. Cholinergic-dependent vasodilation was reduced in MsA from offspring born to obese dams, consistent with previous reports that studied mesenteric arteries from adult offspring in rat maternal obesity models (8, 9, 10). Our results also showed that the maternal obesity-dependent reduction in ACh vasodilation in the offspring is partially mediated by a reduction in endothelial nitric oxide synthesis.

Maternal obesity-dependent reduction in offspring’s MsA ACh vasodilation was prevented by ADN supplementation during late pregnancy in obese dams. This ADN-dependent restoration of cholinergic vasodilation in the offspring vessels seems to be independent of nitric oxide or prostaglandin signaling, since neither L-NAME nor INDO reduced these responses. Another study showing decreased endothelium-dependent vasodilation in rat offspring MsA from high-fat-fed dams have described a compensation by endothelial derived hyperpolarizing factor (EDHF) to restore the reduced cholinergic signaling in the offspring when dams were supplemented with linoleic acid (10). Although we did not directly explore the role of EDHF, we speculate that EDHF is increased in the offspring vessels from ADN-treated obese dams to preserve cholinergic vasodilation.

Maternal ADN supplementation in late pregnant obese dams has multiple beneficial effects in the offspring not only limited to the peripheral vasculature. For example, ADN supplementation the last four days of pregnancy in obese dams prevents fetal overgrowth and improves placental function (15). Moreover, normalization of maternal adiponectin levels in pregnant obese dams prevents liver steatosis and insulin resistance (20), diminishes cardiac hypertrophy, and restores cardiac function in 3-6 month old offspring of obese mouse dams (18). Our results are consistent with these previous studies and support the conclusion that low circulating maternal ADN plays a critical role in mechanistically linking maternal obesity to poor health outcomes in the offspring. Our results also show a cholinergic-specific effect providing mechanistic data on the cause of programming of vascular disease in the offspring of obese mothers.

Our study has some limitations. First, this study was not powered to perform analysis of sex-specific differences and future studies are needed to address this question. However, in previous studies normalization of maternal adiponectin in late pregnant dams prevented adverse cardiometabolic outcomes in 6-9 months old offspring of both sexes (18, 19). Second, we did not include a group of pregnant dams fed control diet that were treated with ADN. However, these dams have normal adiponectin levels and administration of additional adiponectin results in supraphysiological concentrations of the adipokine (22), a condition without clear translational relevance. Indeed, because maternal adiponectin limits fetal growth through its effect on the placenta (23), chronic maternal infusion of adiponectin in normal pregnant mice causes restricted fetal growth (22).

The link between maternal adiponectin levels and placental function, fetal development and long-term cardiometabolic outcomes is mediated by a direct effect of the adipokine on the placenta (23, 24). Thus, activation of maternal and placental ADN signaling, or specifically in the placenta, may represent a promising approach to mitigate the negative cardiovascular effects of maternal obesity on the adult offspring.

## AUTHOR CONTRIBUTIONS

JHD, JAH, ORV and RAL performed the experiments. ORV, LGM, CGJ, FJR, TLP, TJ and RAL conceived the experiments. MM, SF and RAL analyzed the data. MM and RAL wrote the manuscript, MM, JHD, ORV, LGM, CGJ, FJR, TLP, TJ and RAL edited the manuscript. All authors gave final approval of the submitted and published manuscript.

